# Striatal pathology in Spinocerebellar Ataxia Type 1 mice: A comparative study with Huntington’s disease

**DOI:** 10.64898/2025.12.11.693749

**Authors:** Pragya Goel, Praseuth Yang, Lisa Duvick, Orion Rainwater, Shannah Serres, Brennon O’Callaghan, Rocio Gomez-Pastor, Mustafa Mehkary, Terence Gall-Duncan, Peter Langfelder, X. William Yang, Christopher E. Pearson, Patrick E. Rothwell, Harry T. Orr

**Affiliations:** Department of Pathology and Lab Medicine, University of Minnesota - Twin Cities, MN, USA; Institute of Translational Neuroscience, University of Minnesota - Twin Cities, MN, USA; Department of Neuroscience, University of Minnesota - Twin Cities, MN, USA; Center for Neurobehavioral Genetics, The Jane and Terry Semel Institute for Neuroscience and Human Behavior, University of California, Los Angeles, Los Angeles, CA, USA; Department of Psychiatry and Biobehavioral Sciences, David Geffen School of Medicine, University of California, Los Angeles, Los Angeles, CA, USA; Genetics & Genome Biology, The Hospital for Sick Children, Toronto, ON, M5G 0A4, Canada; Department of Molecular Genetics, University of Toronto, Toronto, ON, M5G 1X8, Canada

**Keywords:** Spinocerebellar Ataxia, Huntington’s disease, CAG disorders, striatum, dopamine receptors, Medium Spiny Neurons

## Abstract

Spinocerebellar ataxia type 1 (SCA1) and Huntington’s disease (HD), are motor diseases caused by CAG expansions in *ATXN1* and *HTT*, where SCA1 shows prominent cerebellar neurodegeneration and HD shows prominent striatal neurodegeneration, particularly in the Medium Spiny Neurons (MSNs). Since human and mouse studies demonstrate progressive striatal vulnerability in SCA1, we examined age-dependent molecular, cellular and functional striatal attributes in SCA1 *(f-ATXN1^146Q/2Q^)* knockin mice, by assessing RNA-sequencing, immunohistochemistry and electrophysiology. Striatal mRNAs are downregulated in SCA1 mice, many in common with HD mice, and specificity in MSNs is supported by the rescue of transcriptomic dysregulation with deletion of mutant *Ataxin1* from MSNs. Immunohistochemistry assessed dopamine receptor 1 (D1R) and 2 (D2R) expression in indirect and direct MSNs. In HD mice (*Htt^Q175/Q7^*), expression of both D1R and D2R proteins in MSNs decreased with age in parallel with their RNA levels. In the SCA1 mouse striatum, D1R protein expression decreased with age as seen in murine HD striatum. In contrast, while D2R protein level was decreased similar to D1R protein at 5-weeks of age, by 40-weeks expression of D2R protein recovered to levels recorded in WT mice. Electrophysiological assessment showed a reduction of excitatory synaptic transmission in SCA1 mouse MSNs, indicating functional deficits early in disease. In contrast to cerebellar and many other aspects of SCA1 pathology known to depend on proper nuclear localization of ATXN1 with an expanded polyglutamine, mutating ATXN1’s nuclear localization failed to correct striatal MSN RNA and protein downregulations, indicating a difference in how ATXN1 exerts its pathological effects between the cerebellum and the striatum. Together, these data provide a molecular and cellular basis of striatal pathology in SCA1.

## INTRODUCTION

SCA1 is an inherited neurodegenerative disease with expanded glutamine encoding CAG repeats in the *ATAXIN-1* gene (*ATXN1*) (Orr et al. 1993; Paulson et al. 2017). In SCA1 motor dysfunction is typically associated with dysfunction and degeneration of the cerebellum, in particular Purkinje neurons, a prominent and consistent pathological feature of SCA1 (Zoghbi and Orr 2009; Robitaille, Schut, and Kish 1995). Yet, *ATAXIN-1* is widely expressed throughout the brain (Servadio et al. 1995), raising the possibility that pathology in other brain regions contribute to symptoms seen in SCA1 patients. In Huntington’s Disease (HD), another polyglutamine neurodegenerative disease caused by CAG repeat expansion are in the *Huntingtin (HTT)* gene, motor symptoms result mainly from degeneration of Medium Spiny Neurons (MSNs) within the striatum (Koch et al. 2022; Badreddine et al. 2025; Ferrante, Kowall, and Richardson 1991; Tippett et al. 2017). Importantly, somatic expansion of the *HTT* CAG is considered to be a critical aspect of disease onset (Genetic Modifiers of Huntington’s Disease 2015; Handsaker et al. 2025; Matlik et al. 2024). Curiously, *mATXN1* CAG tract somatic expansions occur in the striatum of SCA1 patients and mouse models similar to the CAG repeat instability seen in HD patients and mouse models (Watase et al. 2003; Mouro Pinto et al. 2020; Chung et al. 1993). As seen in HD, in *f-Atxn1^146Q/2Q^*knockin mice striatal CAG somatic instability occurs in MSNs (Duvick et al. 2024). Additionally, mice in which the *ATXN1^146^* allele is deleted from striatal MSNs show an improvement in rotarod performance at 30-weeks-of-age (Duvick et al. 2024). This latter finding is reminiscent of longitudinal MRI brain volume studies showing significant decrease of striatal volume in SCA1 patients after disease onset (Koscik et al. 2020; Reetz et al. 2013).

Models of basal ganglia function are typically based on the activity of two pathways originating from separate populations of MSNs in the striatum, the Direct pathway from dopamine D1 receptor (D1R) expressing MSNs and the Indirect pathway from dopamine D2 receptor (D2R) expressing MSNs, that connect to output circuits affecting different aspects of motor behavior (Roth and Ding 2024; Kravitz, Tye, and Kreitzer 2012; Sheng et al. 2019; Liang et al. 2022; Durieux, Schiffmann, and de Kerchove d’Exaerde 2012). A central feature of HD neuropathology is the differential vulnerability (cell loss) of MSN subtypes. Indirect pathway (D2R-expressing) MSNs are lost earlier and more extensively than direct pathway (D1R-expressing) MSNs (Reiner et al. 1988; Galvan et al. 2012). The earlier loss of indirect-MSNs is thought to underlie the involuntary movements (chorea) as an early motor symptom in HD (Albin et al. 1990). This early pathological imbalance, which disinhibits the thalamocortical circuits and favors motor output, is thought to be the primary driver of the characteristic choreiform movements of early-to-mid-stage HD (Reiner et al. 1988; Barry et al. 2018; Callahan, Wokosin, and Bevan 2022; Nair et al. 2022). At the molecular level, HD is associated with transcriptional dysregulation of numerous genes, including those encoding the D1R and D2R receptors being progressively downregulated (Burns et al. 2025; Matsushima et al. 2023; Augood, Faull, and Emson 1997). In HD mouse models, similar dysregulated transcriptomic signatures are found in an allelic series of mutant huntingtin knockin mice (Langfelder et al. 2016) and in BAC transgenic mouse model of HD expressing mutant Huntingtin with expanded uninterrupted CAG repeats (Gu et al. 2022).

The parallel emergence of striatal pathology in both SCA1 and HD provides a basis to investigate the fundamental principles governing neuronal vulnerability to polyQ toxicity. Although both diseases are caused by polyQ expansions in ubiquitously expressed proteins, their canonical phenotypes—cerebellar ataxia in SCA1 versus striatal chorea in HD—are distinct. By comparing how two mutant polyQ proteins, Ataxin-1 and Huntingtin, affect the same neural circuits, one can begin to dissect aspects of pathology that are common to a general polyQ insult (e.g., transcriptional repression) and which are specific to the context of the particular protein and its interactome. Currently, a detailed characterization of the molecular, cellular, and functional deficits within the SCA1 striatum is lacking. Key questions that remain unanswered include: (1) What is the precise nature and temporal progression of MSN dysfunction and protein expression changes in SCA1? (2) To what extent does the molecular pathology in the SCA1 striatum, particularly at the level of transcription and dopamine receptor expression, mirror or diverge from that observed in HD? (3) Most importantly, does striatal pathology in SCA1 operate through the same pathogenic mechanisms—specifically, the requirement for nuclear localization of mutant ataxin-1—that are known to drive cerebellar degeneration? To address these questions, we investigated molecular, cellular, and functional pathology of the striatum in SCA1 and HD knock-in mouse models. Using RNA-sequencing, immunohistochemistry, and electrophysiology, we sought to create a diverse profile of striatal deficits as the disease progresses. Together, the data provide a robust cellular and molecular basis for striatal pathology in SCA1, with similarities and differences with HD.

## RESULTS

### Gene expression in D1 and D2 MSNs is reduced in *f-ATXN1^146Q/2Q^* and *Htt^175Q/2Q^* mice

Extensive somatic expansion of the *ATXN1* CAG repeat is found in the striata of SCA1 patients and in knockin SCA1 mice (Watase et al. 2003; Mouro Pinto et al. 2020; Kacher et al. 2024; Zuhlke et al. 1997), similar to the expansion in *Htt* CAGs in HD (Handsaker et al. 2025; Matlik et al. 2024; Kacher et al. 2024; Mouro Pinto et al. 2020). These expansions occur in Medium Spiny Neurons (MSNs) as expansions are significantly reduced when the mutant allele is deleted specifically from MSNs (*f-ATXN1^146Q/2Q^; Rgs9-Cre* mice; (Duvick et al. 2024)). Interestingly, we found that *ATXN1* CAG expansions increase with age in *f-ATXN1^146Q/2Q^* mice (Fig. S1A), corresponding with deficits in motor performance on the rotarod (Fig. S1B; (Duvick et al. 2024)), suggesting that further somatic expansion of the inherited expanded *ATXN1* CAG in the striatum contributes to defects in motor performance as the disease progresses in *f-ATXN1^146Q/2Q^* mice. Of note, the data indicate that once a substantial number of mutant *ATXN1* striatal somatic expansions reach 150 repeats there is a significant impact on motor dysfunction (Fig. S1B), similar to the pathogenic impact 150 repeats in *HTT* have on the striatum of HD patients (Handsaker et al. 2025) and HD mouse models (Wang et al. 2025). We thus sought to characterize molecular, cellular, and electrophysiological attributes in the striata of SCA1 mice.

Previous RNA sequencing shows that the number of differentially expressed genes (DEGs) in a comparison between wildtype (WT) and mutant *Atxn1* mice is the largest in the cerebellum (Handler et al. 2023) while for mutant *Htt* mice compared to WT, this number is the highest in the striatum (Langfelder et al. 2016). An overlap of DEGs was found in striatal DEGs between SCA1 and HD mice. To compare the transcriptome profiles in SCA1 and HD in the cerebellum and striatum, we first correlated Differential Expression (DE) Z statistics for the SCA1 mouse model with expanded repeats in the *Ataxin1* gene vs. WT with DE Z statistics for knockin HD models Q140 and Q175 vs. their controls (Fig. 1A). The latter includes the published data in the striatum of these HD knockin mouse models as well as our new analyses on the cerebellum of these mice (see Methods).

**Figure 1:**
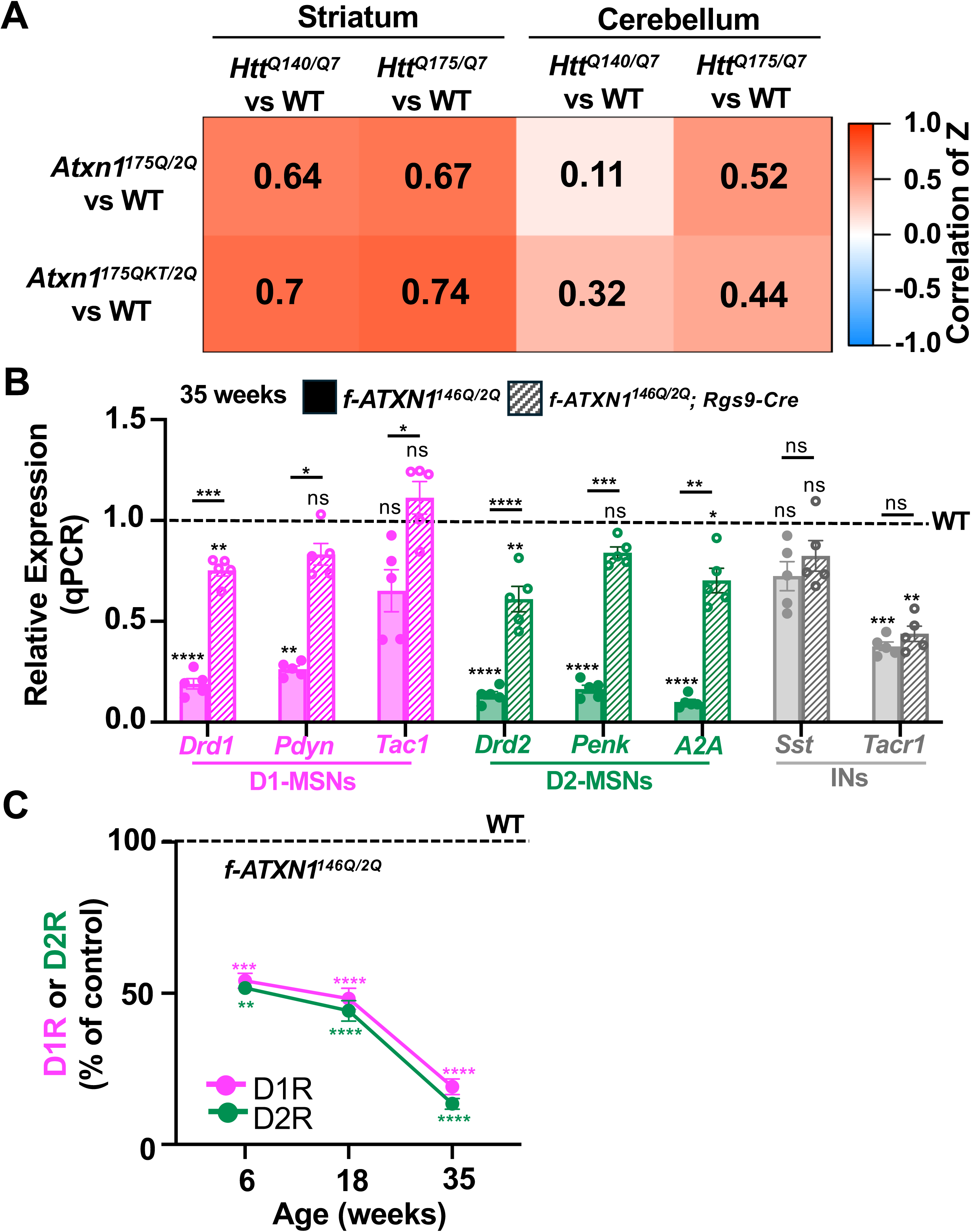
Striatal transcripts are downregulated in SCA1 mice and rescued with deletion of mutant *ataxin1* from MSNs. (**A**) Heatmap representation of pairwise correlations of DE z-statistics across 2 SCA1 (*Atxn1^175Q/2Q^* and *Atxn1^175QK772T/2Q^*), and 2 HD mouse models (*Htt^Q140/Q7^* and *Htt^Q175/Q7^*) in the cerebellum and striatum from 26-week old mice. Wild type (WT) is *Atxn1^2Q/2Q^* for the SCA1 mouse models and *Htt^Q7/Q7^* for the HD mouse models. Correlations larger than 0.2 are shown as numbers in the heatmap. (**B**) Bar plots of gene expression as measured by RNA-seq in *f-ATXN1^146Q/2Q^ and f-ATXN1^146Q/2Q^ X Rgs9-Cre* relative to controls (*Atxn1^2Q/2Q^-*dashed black line) in 35-week old mice. Note the rescue in both D1- and D2-MSNs but not in INs. (**C**) Line graph of gene expression as measured by RT-qPCR for type1 and type 2 dopamine receptors (D1R and D2R) in *f-ATXN1^146Q/2Q^* mice relative to controls (*Atxn1^2Q/2Q^-* dashed black line) at 6, 18, and 35 weeks showing a progressive reduction in mRNA levels of both receptors. Unpaired t test with Welch’s correction; *p=0.05; **p 0.01; ***p=0.001; ****p=0.0001, ns=not significant, p>0.05. See Supplementary Table 1 for complete statistics.

The correlation between Z statistics of *Atxn1^175Q/2Q^*vs WT (*Atxn1^2Q/2Q^*) and *Htt^Q140/Q7^* or *Htt^Q175/Q7^* vs controls (*Htt^Q7/Q7^*and *Htt^Q20/Q7^*) was substantially higher in the striatum compared to the cerebellum. Interestingly, this higher correlation in the striatum compared to the cerebellum persisted when the nuclear localization sequence was mutated in the expanded repeat sequence of *Atxn1* (*Atxn1^175QK772T/2Q^* vs WT), consistent with the observation that preventing nuclear entry of ATXN1 has little effect in the striatum compared with the cerebellum (Handler et al. 2023). The concordance of DE Z statistics between SCA1 and HD mice supports the idea that pathological mechanisms occurring in HD may also drive disease progression in the striatum of SCA1 mice.

To further characterize the striatal transcriptomic profile in SCA1 mice, we analyzed transcript levels of marker/identity genes for specific cell types. MSNs are the main neuron type, constituting 95% of the neurons in the striatum. Substantial pathology has been observed in these striatal cells in HD (Morigaki and Goto 2017; Ehrlich 2012). MSNs are classified as D1- or D2-type (D1-MSNs or D2-MSNs) depending on the expression of type 1 or 2 dopamine receptors (D1R or D2R). D1- and D2-MSNs, respectively, make up the classical direct and indirect pathway with opposing effects on movement, motivation and motor learning (Roth and Ding 2024; Kravitz, Tye, and Kreitzer 2012). The indirect pathway is impacted earlier and more substantially in HD (Reiner et al. 1988; Barry et al. 2018; Nair et al. 2022). RNA qPCR analysis of 35-week-old *f-ATXN1^146Q/2Q^* mice showed significant reductions in transcripts of both D1 and D2-MSN marker genes as well as transcripts in interneurons (INs), the remaining 5% neuron types of the striatum (Fig. 1B). This downregulation is very similar to that observed in *Htt^Q175/Q7^* mice (Langfelder et al. 2016). To assess if this was specific to MSNs, we crossed *f-ATXN1^146Q/2Q^*mice to *Rgs9-Cre* mice that drives selective Cre-recombinase expression in MSNs (Dang et al. 2006; Wang et al. 2014).

In *f-ATXN1^146Q/2Q^; Rgs9-Cre* animals, mRNA levels in D1 and D2-MSNs were restored to nearly WT levels. Importantly, mRNA levels were not restored in INs confirming specificity of Rgs9-Cre in MSNs (Fig. 1B). Finally, given the importance of dopamine receptors (DRs) in motor learning (Sheng et al. 2019; Durieux, Schiffmann, and de Kerchove d’Exaerde 2012) and that they are differentially impacted in HD (Burns et al. 2025; Matsushima et al. 2023; Augood, Faull, and Emson 1997), mRNA levels of D1R and D2R were quantified at 3 different ages – 6-, 24- and 35-weeks to model early, mid and late stage of disease. Both D1R and D2R mRNA levels were reduced by ∼50% as early as 6-weeks-of-age and showed progressive reduction with increasing age (Fig. 1C). Taken together, this transcriptional downregulation data supports the idea of there being similar molecular changes in SCA1 and HD striatum.

### D1R and D2R protein expression in *f-ATXN1^146Q/2Q^* and *Htt^175Q/2Q^* mice

Given the reduction in striatal DR mRNA levels, we assessed expression of the DR proteins. Expression of DRs were quantified in *f-ATXN1^146Q/2Q^* mice and WT controls (*Atxn1^2Q/2Q^*) at 5-, 24- and 40-weeks-of-age in the dorsolateral (DLS) and nucleus accumbens (NAc) subregions of the striatum. These ages were chosen to assess striatal pathology and correlate them with corresponding behavioral data available from previous studies in our lab to monitor motor deficit onset and progression (Duvick et al. 2024; Handler et al. 2023). DLS and NAc striatal subregions were chosen given their role in motor and motivational behaviors (Thorn et al. 2010; Yin et al. 2009; Cataldi et al. 2022). Using Immunofluorescence (IF), *f-ATXN1^146Q/2Q^* and *Htt^175Q/2Q^* mouse striatal coronal sections were stained for D1R and D2R antibodies (Fig 2A). These antibodies were validated using D1R and D2R knockout mice (Uchigashima et al. 2016) to ensure specificity and optimized for immunostaining to yield consistent and reproducible results (Narushima et al. 2006; Uchigashima et al. 2007; Uchigashima et al. 2016). Confocal images were analyzed for D1R or D2R signal intensity using imageJ software. In *f-ATXN1^146Q/2Q^* mice, both DRs showed a significant ∼40% reduction in the DLS at 5-weeks (Fig. 2B). Reduced expression of D1R and D2R proteins were also observed at 5-weeks in the NAc by immunohistochemistry (Fig. S2). At 24-weeks, D1R protein expression continued to decrease (50% reduction) while D2R protein expression showed a recovery compared to its expression at 5-weeks (Only 25% reduction at 24-weeks compared to a 40% reduction at 5-weeks; (Fig. 2B)). At 40-weeks, D1R expression was ∼30% of WT at 40-weeks. In contrast, D2R expression showed a “recovery” in f-*ATXN1^146Q/2Q^* mice with levels nearly equal to wild type, 92% verses 100%; (Fig. 2B)). In *Htt^Q175/Q7^*mice, we observed a progressive trend in reduction of both DRs with D1Rs slightly more reduced than D2Rs at all 3 ages (Fig. 2C).

**Figure 2:**
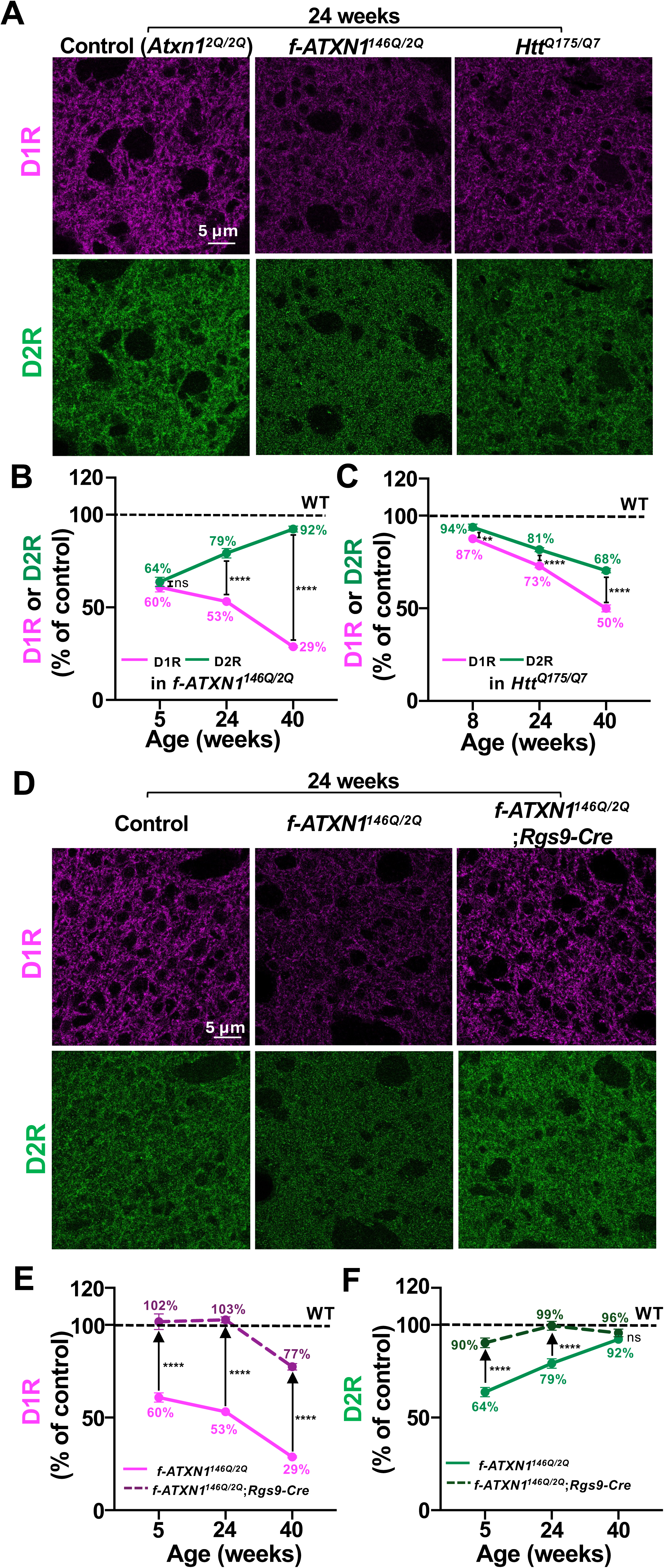
Expression of D1R and D2R proteins with disease progression in SCA1 and HD mice, and correction with MSN deletion of 146Q in SCA1. (**A**) Representative images of single sections of a z-stack acquired using confocal microscopy. Immunofluorescent staining using antibodies against D1R and D2R in the dorsolateral striatum (DLS) of 24-week old *f-ATXN1^146Q/2Q^*(SCA1), *Htt^Q175/Q7^*(HD) and *Atxn1^2Q/2Q^*(control) mice. (**B-C**) Line graphs demonstrating changes in fluorescence intensities of D1Rs and D2Rs in the dorsolateral striatum of *f-ATXN1^146Q/2Q^*(**B**) or *Htt^Q175/Q7^*(**C**) mice relative to respective controls (*Atxn1^2Q/2Q^* (**B**) or *Htt^Q7/Q7^* (**C**): dashed black line at 100%) at 5/8, 24 and 40 weeks. Note the increase in D2R protein levels with age in *f-ATXN1^146Q/2Q^* mice. Significance values are for differences between D1R and D2R changes at each age. (**D**) Representative images of single sections of a z-stack acquired using confocal microscopy. Immunofluorescent staining using antibodies against D1R and D2R in the dorsolateral striatum (DLS) of 24-week old *Atxn1^2Q/2Q^*(control), *f-ATXN1^146Q/2Q^*(SCA1), and *f-ATXN1^146Q/2Q^ ; Rgs9-Cre* (MSN rescue of SCA1) mice. (**E-F**) Line graphs demonstrating a rescue (upward black arrows) in the reduced D1R (**E**) or D2R (**F**) protein fluorescence intensities in the DLS of *f-ATXN1^146Q/2Q^; Rgs9-Cre* mice (MSN rescue of SCA1) shown by darker dashed lines for D1Rs (**E**) and D2Rs (**F**) relative to respective control levels of each receptor type (*Atxn1^2Q/2Q^* ;dashed black line at 100%) at 5, 24 and 40 weeks. Note a significant increase in protein levels with rescue except for D2Rs at 40 weeks where a reduction was not observed (also Fig. 2B). Data is presented from 40-42 images across 3 mice for each genotype and normalized as percentage of average intensity of 40-42 images from 3 control mice. Unpaired t test with Welch’s correction; **p 0.01; ****p=0.0001, ns=not significant, p>0.05. See Supplementary Table 1 for complete statistics.

MSN-specific deletion of *ATXN1^146Q^* using *Rgs9-Cre* restored expression of both DRs to near wild type levels at all 3 ages in the DLS (Fig. 2D-F). Expression of DARPP32 and Tyrosine Hydroxylase (TH) were assessed as readouts of MSN activity and dopamine metabolism in *f-ATXN1^146Q/2Q^* mice. DARPP32 and TH protein levels in the DLS showed a progressive decrease with age from 5- to 24-to 40-weeks (Fig. S3).

### The NLS K772T mutation failed to rescue D1R and D2R protein expression in the striatum of *f-ATXN1^146Q/2Q^* mice

Proper nuclear localization of ATXN1^175Q^ is critical for many of the SCA1-like phenotypes seen in *Atxn1^175Q/2Q^* versus *Atxn1^175QK772T/2Q^* knockin mice (Handler et al. 2023). To investigate the importance of nuclear localization of mutant Atxn1 in the striatum we quantified DR protein levels in *Atxn1^175Q/2Q^* knockin mice in which CRISPR-Cas9 was used to introduce the amino acid alteration (K772T) in the nuclear localization sequence of the expanded ATXN1 protein (*Atxn1^175QK772T/2Q^*; (Handler et al. 2023)). Quantification of DRs in the DLS at all the ages in *Atxn1^175QK772T/2Q^*mice revealed a similar progressive reduction in the protein levels of D1R (Fig. 3B, C) and recovery of D2R (Fig. 3D,E) as in *f-ATXN1^146Q/2Q^* animals (Fig. 2). Transcripts of MSN marker genes also failed to be corrected in the striatum of *Atxn1^175QK772T/2Q^*mice (Fig. 3A). Thus, preventing nuclear localization of mutant ATXN1^175Q^ had no effect on striatal protein level reductions suggesting that Atxn1 exerts its toxic effects in the striatum through a mechanism distinct from other regions of the CNS including the cerebellum.

**Figure 3:**
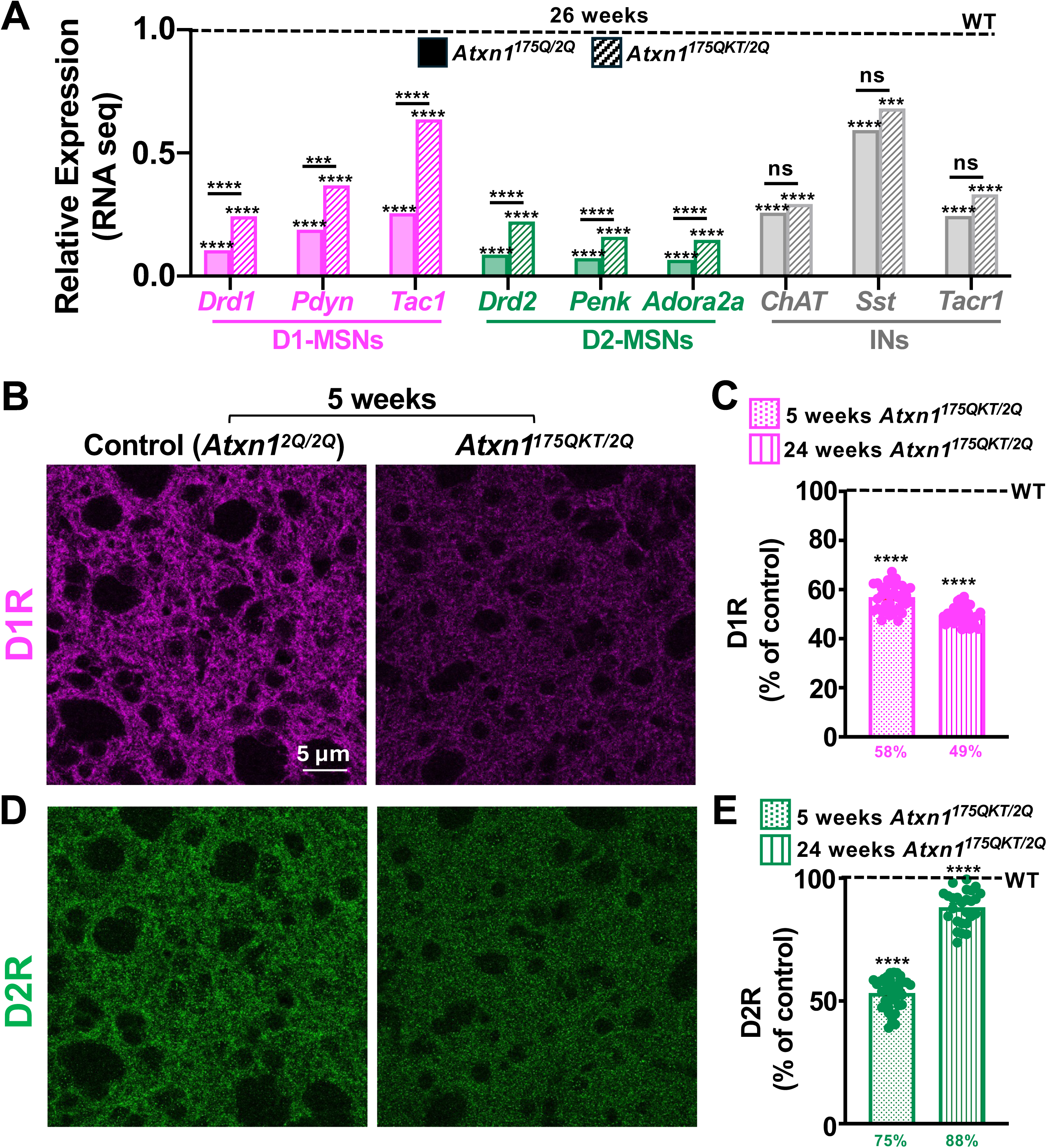
The K772T mutation fails to rescue the downregulation in D1R and D2R protein and MSN transcripts in SCA1 mice. (**A**) Bar plots of reduced gene expression in *Atxn1^175Q/2Q^* (filled bars) and *Atxn1^175QKT/2Q^* (empty bars) relative to controls (*Atxn1^2Q/2Q^* ;dashed black line at 1.0) derived from RNA sequencing data in 26 week old mice for specific marker genes at D1-MSNs (Dopamine receptor type 1(*Drd1*); Prodynorphin (*Pdyn*); Tachykinin Precursor 1 (*Tac1*)), at D2-MSNs (Dopamine receptor type 2(*Drd2*); Proenkephalin (*Penk*); Adora-2A receptor (*A2A*), and at Interneurons (INs; Choline acetyltransferase (*ChAT*); somatostain (*Sst*); Tac1 receptor (*Tacr1*)). Note the lack of rescue in gene expression levels in *Atxn1^175QKT/2Q^* mice. (**B, D**) Representative images of single sections of a z-stack acquired using confocal microscopy. Immunofluorescent staining using antibodies against D1R and D2R in the dorsolateral striatum (DLS) of 5-week old *Atxn1^175QKT/2Q^* and *Atxn1^2Q/2Q^*(control) mice. (**C, E**) Quantification of fluorescence intensities for D1Rs (**C**) and D2Rs (**E**) at 5 weeks or 24 weeks in the DLS of *Atxn1^175QKT/2Q^* mice relative to controls (*Atxn1^2Q/2Q^*; dashed black line at 100%). In (**C)** and **(E)**, data is presented from 40-42 images across 3 mice for each genotype and normalized as percentage of average intensity of 40-42 images from 3 control mice. Unpaired t test with Welch’s correction; ****p=0.0001, ns=not significant, p>0.05. See Supplementary Table 1 for complete statistics.

### Electrophysiological changes in D1 and D2 MSNs in *f-ATXN1^146Q/2Q^* mice

Since the RNA-sequencing data indicated changes in key excitatory pre- and post-synaptic proteins, namely vGlut1, vGlut2, homer1 and PSD95 (Fig. S4), we assessed if excitatory synapse function was impacted in *f-ATXN1^146Q/2Q^* mice. To decouple MSN subtype-specific functional effects, triple transgenic mice containing td-tomato tagged D1-MSNs, eGFP tagged D2-MSNs and the floxed ataxin1 gene with the expanded repeat sequence were generated to identify each MSN subtype and record specifically from D1 or D2-MSNs in SCA1 mice (*f-ATXN1^146Q/2Q^; D1-tdTomato; D2-eGFP*) and controls (*Atxn1^2Q/2Q^; D1-tdTomato; D2-eGFP*). Using whole-cell voltage-clamp electrophysiology, we recorded miniature excitatory postsynaptic currents (mEPSCs) at D1 and D2 MSNs in 5-8 week animals. Although mEPSC frequency was reduced at both D1-MSNs and D2-MSNs (Fig. 4B,C left columns), we found no differences in the mEPSC amplitude in either MSN subtype (Fig. 4B,C right columns). This was surprising given the ∼50% reduction in excitatory postsynaptic mRNA expression levels (Fig. S4). The reduction in mEPSC frequency could be due to a reduction in the number of synapses, a reduction in presynaptic release, a change in the number of dendritic spines, or a combination of these possibilities. This functional result differs from that observed in a study using the *Htt^Q175/Q7^* HD mouse model where a reduction in mEPSC frequency was found only in D1-MSNs, and shown to likely be due to altered dendritic topology and reduced spine density specifically on D1-MSNs (Goodliffe et al. 2018). However, other studies using BACHD and YAC128 mouse models have reported an mEPSC reduction and changes in dendritic morphology at both MSN subtypes (Andre, Fisher, and Levine 2011; Park et al. 2025), broadly suggesting functional changes at both MSN subtypes in HD mouse models. Similar to *f-ATXN1^146Q/2Q^* mice, no significant differences in mEPSC amplitude were observed in HD mice in either MSN subtype (Goodliffe et al. 2018; Andre, Fisher, and Levine 2011; Indersmitten et al. 2015). Together, this suggests that functional deficits start to manifest in both MSN subtypes at an early time point in *f-ATXN1^146Q/2Q^* mice when both DR types are similarly reduced (Fig. 2A). Thus, both upstream and downstream pathways of MSNs in SCA1 mice are likely impacted early in disease.

**Figure 4:**
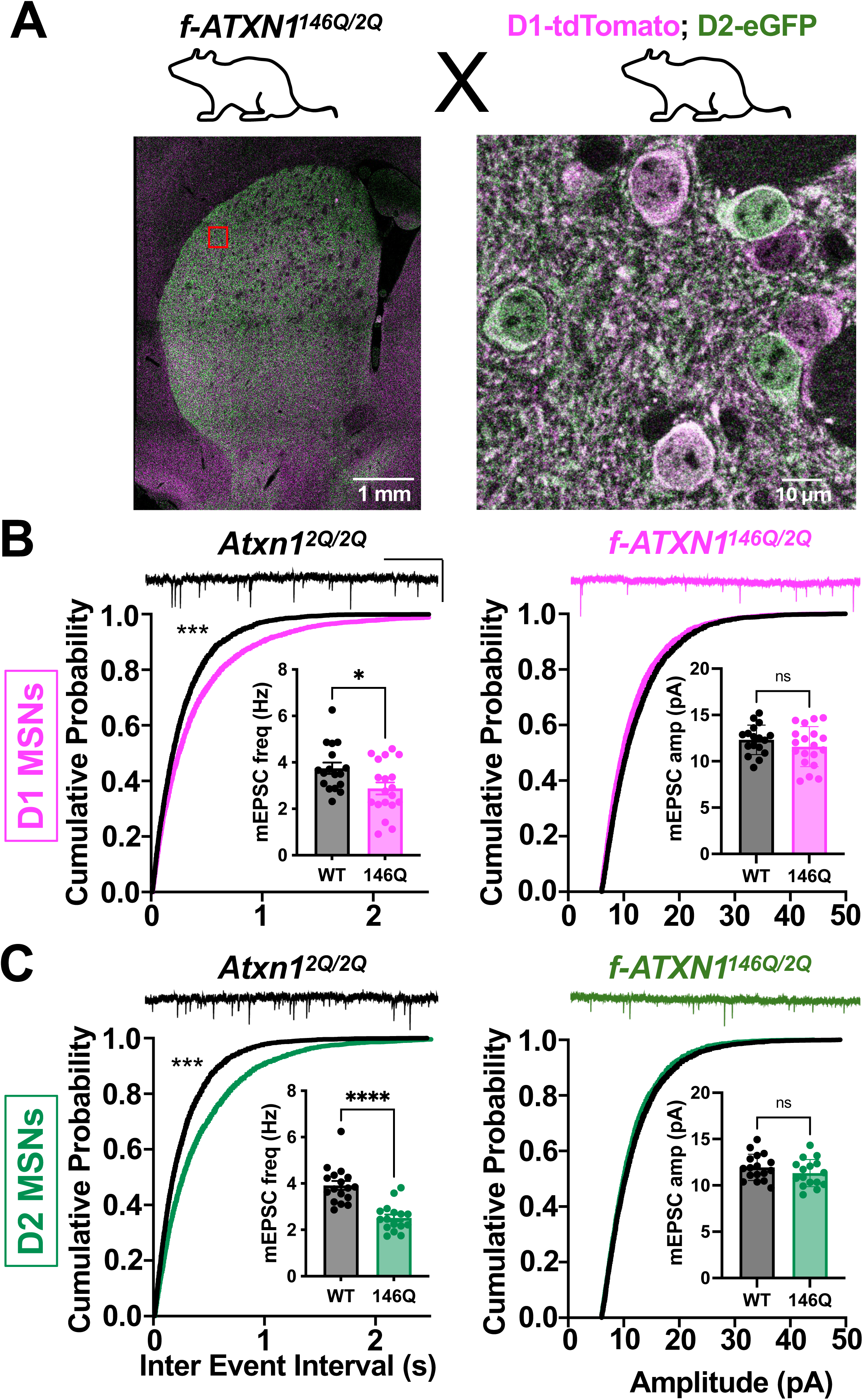
Reduced excitatory current frequency on D1- and D2-MSNs in SCA1 mice. (**A**) *f-ATXN1^146Q/2Q^* mice were crossed to double transgenic mice (D1 tagged with tdTomato; D2 tagged with eGFP) to generate triple transgenic mice (*f-ATXN1^146Q/2Q^*; D1-tdTomato; D2-eGFP) and littermate controls (*Atxn1^2Q/2Q^*; D1-tdTomato; D2-eGFP) for D1 (red) or D2-MSN (green) identification. Representative image of the entire striatum (left) with tagged MSNs and stained with a DARPP32 antibody (white) to label all MSNs. A zoomed in image (right). (**B-C**) **Top**, example traces of mEPSCs at D1-MSNs (**B**) or D2-MSNs (**C**) recorded from 5-8 week old SCA1 and control mice. Scale bars: X-axis 2 sec, Y axis 20 pA. **Bottom**, cumulative fraction plots of inter-event interval (IEI; left) and amplitude (right) of mEPSCs for both genotypes. Insets represent average data for mEPSC frequency (left; calculated as inverse of IEI) and mEPSC amplitude (right). SCA1 mice show a decrease in mEPSC frequency in both MSN subtypes ((**B**)-red; D1-MSNs, n=16, (**C**)-green; D2-MSNs, n=16) but no change in mEPSC amplitude compared to control mice (black, n=16). *p=0.05; **p 0.01; ***p=0.001; ****p=0.0001, ns=not significant, p>0.05. Unpaired t test with Welch’s correction for averages, Mann Whitney test for frequency distributions. See Supplementary Table 1 for complete statistics.

## DISCUSSION

HD and SCA1 are polyglutamine neurodegenerative diseases where the expanded CAG repeats are located in two widely expressed genes. Yet, each disease is characterized by distinct clinical characteristics and predominant sites of neuropathology. In HD patients MSNs of the striatum are the most susceptible to degeneration. In SCA1, cerebellar Purkinje cell degeneration is a frequent and prominent pathological feature. However, in SCA1 patients imaging studies show that volumetric changes in the striatum are associated with progression of disease and motor dysfunction (Reetz et al. 2013; Koscik et al. 2020). Data presented here provides a multi-level characterization of striatal molecular and electrophysiological alterations in MSNs with mutant ATXN1 in *ATXN1* knockin SCA1 mouse models; *Atxn1^175Q/2Q^* and *f-ATXN1^146Q/2Q^*. Among the key striatal cellular marker genes whose expressions were downregulated in the SCA1 mice are the genes encoding D1R and D2R, markers of the direct and indirect pathways respectively. While D1R and D2R RNA levels both progressively decrease with age, at the protein level, D1R expression decreased with age while D2R protein level was decreased similar to D1R protein at 5-weeks-of-age but by 40-weeks-of-age D2R protein recovered to essentially that seen in WT mice. The functional impact of the recovery of D2R protein expression is yet to be determined. In contrast to Purkinje neurons of the cerebellum and cells within other regions where the K772T mutation in the ATXN1 NLS disrupts entry of mutant ATXN1 into the nuclei of cells and decreases pathogenesis (Handler et al. 2023; Duvick et al. 2024), the K772T mutation in mutant ATXN1 had no impact on striatal pathogenesis in *f-ATXN1^146Q/2Q^* mice. This intriguing finding indicates that the SCA1 molecular pathogenic pathway in the striatum differs from that in the cerebellum. Electrophysiological assessment showed a reduction of mEPSC frequency in D1 and D2 MSNs indicating that along with the molecular alterations striatal functional deficits are a component of disease.

### Shared Striatal Transcriptional Alterations in SCA1 and HD mice

Data presented in this study position the striatum as a site of convergent pathology between SCA1 and HD mice. In addition to the previously reported progressive somatic repeat instability observed in the striatum in SCA1 mice (Fig. S1; (Duvick et al. 2024)) and HD mice (Wang et al. 2025), the correlation analysis revealed that transcriptional changes in the striatum of SCA1 and HD mice are substantially more similar than those in the cerebellum (Fig. 1A). This suggests that despite being initiated by two different mutant proteins, the downstream transcriptional response to polyglutamine expansion within striatal neurons follows a similar trajectory.

This shared response is characterized by an early and sustained downregulation of genes critical for neuronal identity and function. In *f-ATXN1^146Q/2Q^*mice, transcripts for markers of both D1- and D2-MSNs, as well as interneurons, were significantly reduced by 26-weeks. This process begins much earlier, with downregulation evident at 8-weeks of age, mirroring the timeline in HD mouse models (Garland et al. 2018; Smith et al. 2023; Menalled et al. 2012; Carter et al. 1999; Langfelder et al. 2016) and correlating with the onset of motor deficits.

The transcriptional alterations are cell-autonomous to MSNs, as demonstrated by the recovery of D1R and D2R mRNA levels upon MSN-specific deletion of mutant *ATXN1*. Importantly, transcriptional recovery was not seen in striatal interneurons.

### Immunohistochemistry Reveals Recovery of D2R Protein Expression Specific to MSNs of *f-ATXN1^146Q/2Q^* mice

Despite the striking similarities at the mRNA level, the IF analysis uncovered a profound divergence in the post-transcriptional handling of dopamine receptors, distinguishing *f-ATXN1^146Q/2Q^* mice pathology from that of *Htt^175Q/Q7^* mice. In SCA1 mice, both D1R and D2R protein levels were significantly reduced in the dorsolateral striatum at 5-weeks of age. However, as the disease progressed, their expression patterns diverged dramatically: D1R protein levels continued to decline through 40-weeks, while D2R protein underwent a remarkable “recovery,” returning to near-wildtype levels. This is in stark contrast to HD mice, where both D1R and D2R proteins showed a progressive reduction with age (Fig. 2A-C).

This recovery of D2R protein in the face of persistently low levels of its corresponding mRNA implies activation of a, SCA1-specific post-transcriptional compensatory mechanism. In the SCA1 context, this may be due to increased D2R protein translation, decreased D2R protein degradation, altered protein trafficking and/or altered alternative splicing, or an upregulation of D2Rs in the axonal terminals of dopaminergic neurons (i.e. D2 autoreceptors). This is an important finding as it demonstrates that transcriptional profiles do not always predict the proteomic landscape in neurodegenerative disease and reveals a previously unknown cellular response unique to mutant ataxin-1. A similar difference in D2R mRNA and protein levels was reported in the MPTP Parkinson’s model (Todd et al. 1996), and this mismatch in mRNA and protein expression has been noted in aging and neurodegenerative diseases specifically Alzheimer’s, Parkinson’s and Huntington’s disease (Buck et al. 2025; Ramazi et al. 2024).

### Striatal Pathogenesis is Independent of Ataxin-1 Nuclear Localization: A mechanistic distinction from the cerebellum

An intriguing finding of this study is that the pathogenic mechanism driving striatal dysfunction in *f-ATXN1^146Q/2Q^* mice is independent of mutant Ataxin-1’s nuclear localization. In the cerebellum, the nuclear entry of Ataxin-1 is an absolute prerequisite for its toxicity (Handler et al. 2023). This is evident by the 87% correction of DEGs in the cerebellum but only a 53% correction in the striatum of *Atxn1^175QK772T/2Q^* compared to *Atxn1^175Q/2Q^*mice (Handler et al. 2023). Further, the number of ATXN1 nuclear inclusions is significantly reduced in cerebellar Purkinje cells with the K772T mutation (*Atxn1^175QK772T/2Q^*), is reduced in the 5-week striatum but is similar to inclusions in the 12-week striatum in *Atxn1^175Q/2Q^* mice. This suggests a somatic expansion dependent, secondary nuclear entry pathway for striatal MSNs but the cerebellar Purkinje cells in SCA1 mice. Here, we show that introducing the K772T mutation, which alters Ataxin-1’s nuclear import, failed to prevent either the transcriptional downregulation of a selection of MSN marker genes (specific marker genes at D1-MSNs (Dopamine receptor type 1(Drd1); Prodynorphin (Pdyn); Tachykinin Precursor 1 (Tac1)), at D2-MSNs (Dopamine receptor type 2 (Drd2); Proenkephalin (Penk); Adora-2A receptor (A2A), and at Interneurons (INs; Choline acetyltransferase (ChAT); somatostain (Sst); Tac1 receptor (Tacr1)) or the dynamic changes in D1R and D2R protein levels in the striatum of *Atxn1^175QK772T/2Q^* mice (Fig. 3). This result was consistent with our transcriptomic analysis, which showed that the high correlation in striatal DEGs between SCA1 and HD persisted even with the NLS mutation (Fig. 1A). This finding demonstrates a fundamental mechanistic bifurcation in SCA1 pathogenesis: the striatum and cerebellum are functionally vulnerable to mutant Ataxin-1 through distinct pathways and highlights the critical importance of cell context in shaping the mechanisms of polyglutamine neurodegeneration.

### Distinct Striatal Functional Deficits in SCA1 and HD mice Indicate Differential Vulnerability of Striatal Circuits

While further work is needed to understand how molecular changes observed manifest in functional deficits, the loss of D1R protein expression along with the recovery of D2R expression suggests a selective and progressive vulnerability of the direct pathway in *f-ATXN1^146Q/2Q^* mice. This contrasts with observations in HD mice, where the indirect (D2R-expressing) pathway is considered more susceptible transcriptionally (Matsushima et al. 2023), although it is electrophysiologically unaffected in HD knockin mouse models (Goodliffe et al. 2018; Andre, Fisher, and Levine 2011) .

The electrophysiological results suggest certain fuctional differences in the two diseases. In early-stage *f-ATXN1^146Q/2Q^* mice, the frequency of miniature excitatory postsynaptic currents (mEPSCs) was significantly reduced in *both* D1- and D2-MSNs. This indicates a deficit in excitatory synapse numbers and/or presynaptic glutamate release affecting both the direct and indirect pathways. This finding is different from that seen in one study in a Q175 HD mouse model where D1 and D2-MSNs were tagged, and a reduction in mEPSC frequency was found to be specific to D1-MSNs, and was attributed to a reduction in dendritic spines specifically on D1 and not D2-MSNs (Goodliffe et al. 2018). Another study using the BACHD and YAC128 mouse models reported reduced mEPSCs in both D1- and D2-MSNs although just 1 MSN subtype was tagged and the other subtype was assumed to be the correct one (Andre, Fisher, and Levine 2011). Dendritic morphological changes have been shown to occur in both D1 and D2 MSNs in HD Q140 knock in mice (Park et al. 2025). In untagged HD animals where D1 and D2-MSNs were indistinguishable, an overall reduction in mEPSC frequency (Indersmitten et al. 2015) and dendritic spines (Indersmitten et al. 2015; Klapstein et al. 2001) was reported. These early and widespread synaptic deficits indicate that, at a circuit level, mutant Ataxin-1 and Huntingtin may disrupt certain aspects of striatal function (miniature excitatory current frequencies) at both MSN subtypes. To parse out specific differences between SCA1 and HD at a functional level, further electrophysiological characterization with a direct comparison between the two models is needed in the future.

### Conclusion and Future Implications

In summary, this study further identifies the striatum as a key locus of pathology in SCA1 knockin mice. This work demonstrates that while SCA1 and HD mouse models share a common vulnerability to transcriptional dysregulation in MSNs, they diverge in MSN subtype vulnerability in terms of protein expression and function. HD is also distinct from SCA1, in that the former shows neurodegenerative vulnerability in the striatum, while the latter shows neurodegenerative vulnerability in the cerebellum. The unique recovery of D2R protein in *f-ATXN1^146Q/2Q^* mice, the impact on excitatory synaptic function, and, most importantly, the discovery of a nuclear-localization-independent toxic mechanism collectively advance our understanding of how mutant Ataxin-1 affects the striatum.

Areas of future focus include elucidating the specific post-transcriptional mechanisms driving D2R recovery in *f-ATXN1^146Q/2Q^*mice and identifying the pathways responsible for the NLS-independent toxicity of Ataxin-1 in MSNs. Understanding these cell-type-specific pathogenic processes will be essential for developing targeted therapies that can address the full spectrum of symptoms in SCA1 and other polyglutamine disorders.

## MATERIALS AND METHODS

### Mice

Experiments were performed with female and male mice maintained on a C57BL/6J genetic background.

Mice were kept on a 12:12 day/night cycle at 23 degrees Celsius with food and water provided *ad libitum*. In all experiments, nearly equal numbers of male and female mice were used. Mice were age-matched within experiments and corresponding littermate controls were used. All mice were housed and managed by Research Animal Resources under specific pathogen-free conditions in an Association for Assessment and Accreditation of Laboratory Animal Care International approved facility. Experimental procedures were approved by the Institutional Animal Care and Use Committee of the University of Minnesota.

*f-ATXN1^146Q/2Q^* conditional knock-in mice were generated as described (Duvick et al. 2024). Briefly, coding exons 7-8 with the intervening intron of one allele of the mouse *Atxn1* gene was replaced with the human *ATXN1* coding exons 7-8 and the intervening intron using site-specific recombination at flanking FRT and LoxN recombination sites. HD knock-in mice (*Htt^zQ175/Q7^*) were used for IHC experiments and were generously shared by Dr. Rocio Gomez-Pastor at the University of Minnesota. Here the mouse exon 1 of *Htt* is replaced with an human exon 1 of *Htt* harboring a ∼190 CAG repeat expansion (Menalled et al. 2012). The two BAC transgenic reporter mouse lines: D1-tdTomato (The Jackson Laboratory, stock #016204) and D2-eGFP (Mutant Mouse Resource & Research Centers #036931-UCD) were crossed to *f-ATXN1^146Q/2Q^* mice to obtain hemizygous offspring expressing tdTomato in D1-MSNs and/or eGFP in D2-MSNs with or without the 146 CAG expanded repeat in *Atxn1*. *Rgs9-Cre* mice were purchased from The Jackson Laboratory (RRID:IMSR_JAX:020550). Both *Atxn1^175Q/2Q^* knock-in and NLS-defective *Atxn1^175QK722T/2Q^* mice have been described (Handler et al. 2023).

### Genotyping

PCR was performed with the following primers (Integrated DNA Technologies) to determine which animals have an expanded *f-ATXN1^146Q/2Q^* allele: ATXN1-146Q repeat Forward (5’-CAACATGGGCAGTCTGAG) and ATXN1-146Q repeat Reverse (5’-GTGTGTGGGATCATCGTCTG). To assess recombination by Cre the following primer set was used: Recombination Forward (5’-GGGAATGGTACCAACCTT TCTG) and Recombination Reverse (5’-GTAGAACCCCAGACCCTCGT). Cre F oIMR 1084 (5’-GCGGTCTGGCAGTAAAAACTATC) and Cre R oIMR 1085 (5’-GTGAAACAGCATTGCTGTCACTT) were used to genotype all Cre lines. To determine the presence of the D1-tdTomato or D2-eGFP tags, the following primers were used: For D1-tdTomato, BGH-F (CTT CTG AGG CGG AAA GAA CC) and dDR4 (TTT CTG ATT GAG AGC ATT CG); for D2-eGFP, D2 (CCC GAA GCT TCT CGA GGC GCG CCC TGT GCG TCA GCA TTT GGA GCA AC) and e-GFP (TCA GGG TCA GCT TGC CGT AGG).

### Repeat length analysis

Fragment analysis of f-Q146 Atxn1^Q146/Q2^ mouse samples, PCR amplification was performed using the Platinum SuperFi II Mastermix system. For *ATXN1,* 10 ul of 2x SuperFi II Mastermix, 1 ul of 2.5uM forward primer (FAM-5′-CCAGACGCCGGGACACAAGGCTGAG-3’), 1 ul of 2.5 reverse primer (5’-CCGGAGCCCTGCTGAGGTG-3’) with a final reaction volume of 20ul made up with ultrapure water. PCR Cycling conditions were as follows : 98°C 5 min, 30 cycles of 98°C 10 s, 66°C 10 s, 72°C 30 s, followed by a final extension step of 72°C for 5 min. All PCR products were denatured with HiDi formamide and boiling at 95°C for 5 minutes, and then processed by capillary gel electrophoresis with the Genescan 1200LIZ size markers on a 3130xl Genetic Analyzer (Applied Biosystems). Peak Scanner 2 software was used to visualize the repeat sizes, and repeat lengths were calculated by subtracting the length of non-repeat sequence in the PCR product and dividing by 3. Instability indices from GeneMapper peak height data were calculated as previously described ^8^ with a 20% relative height threshold.

### RT-qPCR

Animal tissues were harvested and stored in Invitrogen RNAlater stabilization solution overnight at 4℃ and then stored at -80℃ until use. RNA was extracted from striatum using TRIzol from Invitrogen following the provided protocol and adjusting for total tissue mass. Synthesis of cDNA was performed in duplicate using iScript Advanced cDNA Synthesis kit from Bio-Rad. All RT-qPCR of the genes measured was performed using IDT Predesigned Probe Assays listed in Appendix Table 3. IDT predesigned probe assays are designed to be run on standard cycling conditions with amplification temp of 60℃. *Gapdh* was used as a reference control by which gene expression values were normalized over. Statistical testing was performed using two-sided Student’s *t* test or by one-way ANOVA on each individual gene. Statistically significant ANOVAs were followed up with Tukey’s post hoc tests.

### Differential Expression Analyses Preprocessing

Previously published differential expression analysis results were obtained from supplemental files from HD (Langfelder et al. 2016) and SCA1 (Handler et al. 2023). The HD allelic series utilized DESeq2 R package while the SCA1 nuclear localization study utilized edgeR R package. The HD allelic series provided Entrez IDs and MGI gene symbols while the SCA1 study used Ensembl IDs and newer MGI gene symbols. Matching of appropriate genes was done by using the gconvert R function from the gprofiler2 package to convert HD Entrez IDs to Ensembl IDs or old MGI gene symbols to concurrent Ensembl IDs. Lists were then filtered for genes present in both analyses.

### Immunofluorescence

Mice were deeply anesthetized with Ketamine, transcardially exsanguinated with PBS, and perfused using 10% formalin. Brains were post-fixed overnight in 10% formalin then stored in PBS at 4C until sectioning. Dissected out brains were dehydrated in 30% sucrose + 0.1% sodium azide in PBS overnight or until they sank to the bottom of the tube. Coronal sections of 30μm were cut using a Leica VT 1000S vibratome in ice-cold PBS. Sections were blocked for 1 hour in 10% normal donkey serum and 0.25% Triton X-100 in PBS. Subsequent staining was carried out in 1% normal donkey serum and 0.25% Triton X-100 in PBS (PBST). Sections were incubated for 24-48 hours with primary antibodies at 4C. Following incubation, sections were washed three times in PBS for 30 mins and exposed to secondary antibodies for 2 hours at room temperature. Sections were washed thrice in PBST each time for 10 mins and mounted using ProLong Gold antifade reagent (Thermo Fisher Scientific). The following primary antibodies were used: 1:400 anti-goat DARPP32 (Nittobo Medical; RRID AB_2571684), 1:200 anti-rabbit D2R (Nittobo Medical; RRID AB_2571596), 1:400 anti-guinea pig D1R (Nittobo Medical; RRID AB_2571595), and 1:1000 anti-rabbit Tyrosine Hydroxylase (Millipore Sigma; RRID AB_390204). All secondary antibodies were obtained from Jackson Immunoresearch Labs (Donkey anti-goat Alexa 647; code: 705-605-147, Donkey anti-Rabbit Cy3; code: 711-165-152, Donkey anti-Guniea Pig Alexa 488; code: 706-545-148), reconstituted according to product specifications and used at 1:1000.

### Imaging and analysis

Samples were imaged using a Leica SP8 Confocal microscope equipped with a 63x APO 1.4NA oil immersion objective using separate channels with four laser lines (405 nm, 488 nm, 555 nm, and 647 nm). For fluorescence intensity quantifications of D1R, D2R, DARPP32 and TH, z-stacks were obtained using identical gain and laser power settings with z-axis spacing of 0.3 µm for all genotypes within an individual experiment. Raw confocal image stacks were analyzed. The mean intensities for D1R, D2R, TH and DARPP32 were calculated in imageJ after identical background subtraction using a Rolling Ball Radius of 50 pixels for. All measurements based on confocal images were taken from ∼40-42 images from striatal subregions from 3 animals per genotype.

### Electrophysiology

Mice were deeply anesthetized with isoflurane and perfused with 10 mL ice-cold sucrose cutting solution containing (in mM): 228 sucrose, 26 NaHCO3, 11 glucose, 2.5 KCl, 1 NaH2PO4-H2O, 7 MgSO4-7H2O, 0.5 CaCl2-2H2O. Mice were subsequently decapitated and brains were quickly removed then placed in ice-cold sucrose cutting solution. Coronal slices (240 µm thick) containing nucleus accumbens (NAc) were collected using a vibratome (Leica VT1000S) and allowed to recover submerged in a holding chamber with artificial cerebral spinal fluid (aCSF) containing (in mM): 119 NaCl, 26 NaHCO3, 11 glucose, 2.5 KCl, 1 NaH2PO4-H2O, 2.5 CaCl2-2H2O, 1.3 MgSO4-7H2O. Slices recovered in warm aCSF (33°C) for 15 min and then equilibrated to room temperature for at least one hour before use. Slices were transferred to a submerged recording chamber and continuously perfused with aCSF at a rate of 2 mL/min at room temperature. All solutions were continuously oxygenated (95% O2/5% CO2).

Whole-cell voltage-clamp recordings from MSNs in the NAc core were obtained under visual control using IR-DIC optics from an Olympus BX51W1 microscope. D1-MSNs were distinguished from D2-MSNs by the expression of tdTomato or eGFP, respectively, and held at -70 mV. Borosilicate glass electrodes (3-5 MΩ) were filled with (in mM): 120 CsMeSO4, 15 CsCl, 10 TEA-Cl, 8 NaCl, 10 HEPES, 5 QX-314, 4 ATP-Mg, 1 EGTA, 0.3 GTP-Na (pH 7.2-7.3). Excitatory synaptic transmission was pharmacologically isolated using GABAA receptor antagonist picrotoxin (50µM, Tocris). Miniature EPSCs (mEPSC) were obtained in the presence of tetrodotoxin (500 nM, Fischer Scientific) to block spontaneous activity.

All recordings were performed using a MultiClamp 700B (Molecular Devices), filtered at 2 kHz, and digitized at 10 kHz. Data acquisition and analysis were performed online using Axograph software. Series resistance was monitored continuously and experiments were discarded if resistance changed by >20%.

At least 200 events per genotype were acquired across 18 sec sweeps, filtered at 0.5 kHz, and detected using an amplitude threshold of 6 pA and a signal-to-noise ratio threshold of 4 standard deviations.

### Pairwise correlation analysis (Fig. 1A)

#### Bulk RNA-seq data preprocessing for SCA1 data (Fig. 1A)

We downloaded raw counts from GSE218303. Data from each brain region and age were analyzed as a separate data set. In each data set, we retained only genes with at least 0.3 robust counts per million reads (CPM) in at least the number of samples in the genotype group, generally 4. The rationale for is to only include genes that are likely to be expressed in at least one genotype. The robust CPM is calculated by first normalizing the counts using the Relative Log Expression (RLE) normalization implemented in DESeq2 (Love, Huber, and Anders 2014) and then dividing the normalized counts by the median normalized library size and multiplying by 10^6^. The numbers of retained genes ranged from 18,134 (10w cerebellum) to 19,337 (26w hippocampus).To identify potential outliers, we used a modified version of the sample network methodology originally described (Oldham, Langfelder, and Horvath 2012). Specifically, to quantify inter-sample connectivity, we first transformed the raw counts using variance stabilization (DESeq2 function varianceStabilizingTransformation which also includes RLE normalization) and then used Euclidean inter-sample distance based on the scaled profiles of the 8000 genes with highest mean expression. The intersample connectivities k were transformed to Z scores using robust standardization,

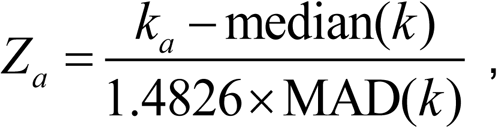

where index a labels samples, MAD is the median absolute deviation, a robust analog of standard deviation, and the constant 1.4826 ensures asymptotic consistency (approximate equality of MAD and standard deviation for large, normally distributed samples). Finally, samples with Z_a_ < -6 were removed. This procedure resulted in the removal of 3 samples from cerebellum at 26-weeks but no samples from any of the other region/age combinations.

#### Bulk RNA-seq data preprocessing for HD mouse model data (Fig. 1A)

The work (Langfelder et al. 2016) reported results of DE analysis in striatum but not in cerebellum. As both striatum and cerebellum RNA-seq data are available from the Gene Expression Omnibus (GEO; accessions GSE65774 and GSE73468, respectively), we downloaded both data sets and aligned them to mm10 genome using STAR followed by HTSeq to summarize the aligned reads to gene counts. We followed an older version of the same preprocessing pipeline as for the SCA1 data that differed from the SCA1 in the following modifications: the low expression threshold was set at 0.8 “plain”, i.e., not robust, CPM and the outlier removal threshold was -4.5, resulting in a more conservative resulting set of DE statistics.

#### Differential expression testing (Fig. 1A)

To make differential expression (DE) testing robust against potential outlier measurements (counts) that may remain even after outlier sample removal, we calculated individual observation weights designed to downweigh potential outliers. The weights are constructed separately for each gene. First, Tukey bi-square-like weights(Wilcox 2012) *λ* are calculated for each (variance-stabilized) observation *x_a_* (index *a* labels samples) as

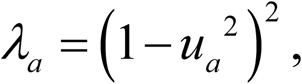

where

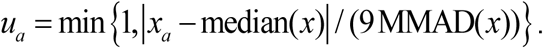

The median is calculated separately for each gene across all samples. MMAD stands for modified MAD, calculated as follows. For each gene, we first set MMAD = MAD. The following conditions are then checked separately for each gene: (1) 10^th^ percentile of the weights *λ* is at least 0.1 (that is, the proportion of observations with weights <0.1 is less than 10%) (Langfelder and Horvath 2012) and (2) for each individual genotype, 40^th^ percentile of the weights *λ* is at least 0.9 (that is, at least 60% of the observation have a weight ≥ 0.9). If both conditions are met, MMAD = MAD. If either condition is not met, MMAD equals the lowest value for which both conditions are met. The rationale is to exclude outliers but ensure that the number of outliers is not too large either overall or in each genotype group. This approach has previously been used in (Lee et al. 2018; Wang et al. 2022).

DE analysis was carried out using DESeq2 (Love, Huber, and Anders 2014) version 1.36.0 (SCA1 model data) amd 1.20.0 (HD model data) with default arguments except for disabling outlier replacement (since we use weights to downweigh potential outliers) and independent filtering (since we have pre-filtered genes based on expression levels). For the HD model data, the control group contained both Q20 and WT (Q7) mice. Sex (both striatum and cerebellum) and batch (cerebellum only as all striatum samples were sequenced in a single batch) were used as covariates in DE testing.

To evaluate transcriptome-wide concordance of effects, we use correlations of Z statistics; the Z statistics are defined as the log_2_ fold changes (technically, the regression model coefficients) divided by their standard errors. Specifically, given two different differential expression analyses (e.g., conditions arranged into two pairs, with DE analysis between the two conditions in each pair), we arrange the Z statistics of genes common to both comparisons as two linear vectors and correlate them. We find correlations of Z statistics generally more consistent and informative than correlations of log fold changes which tend to be influenced by low-expressed genes that often exhibit high but highly variable (i.e., noisy and less statistically significant) log fold changes. A disadvantage of transcriptome-wide correlations is that their statistical significance is not straightforwardly assessed: expression profiles of genes tend to exhibit strong correlation patterns that make naïve significance estimates hopelessly inflated. In our experience, transcriptome-wide correlations of Z statistics around ±0.2 correspond approximately to (one-sided) p-values around 0.05.

### Statistical Analysis

All other statistical tests were performed using GraphPad Prism software (v9.0). All data were checked for normality and met assumptions for using parametric statistical tests. For multigroup comparisons, one-way ANOVAs were performed. When ANOVA findings were significant, (p<0.05), the analysis was followed by multiple comparisons testing using Tukey’s (all pairwise comparisons) correction for multiple comparisons. For Fig. 1A, enrichment was evaluated using the hypergeometric test (also known as Fisher’s exact test) with the set of analyzed genes (after filtering out low-expressed ones) used as the background. All data presented are shown as mean ± SEM. Significant results are denoted as * (p<0.05), ** (p<0.01), *** (p<0.001), and **** (p<0.0001). Complete statistical analysis details for all data analyzed in Prism are presented in Table S1.

## ACKNOWLEDGEMENTS

This study was supported by NIH grants R01NS022920 (H.T.O.), R35NS127248 (H.T.O.). X.W.Y. and P.L. were supported by NINDS/NIH grants (R01NS113612) and CHDI Foundation, Inc. The authors thank Jessica Swanson from the Rothwell Laboratory for demonstrating whole-cell patch clamp electrophysiology to Pragya Goel in the early phases of recordings.

## AUTHOR CONTRIBUTIONS

P.G. and H.T.O conceived the study and wrote the paper. P.G, P.Y and H.T.O designed experiments and interpreted the data. P.G. performed the electrophysiology, microscopy, imaging and related analysis. P.Y conducted the RNA seq and qPCR experiments and analysis. O.R and S.S bred mice and managed the colony. L.D conducted the genotyping. P.L performed the transcriptomic cross-correlation analysis for the striatum verses the cerebellum in SCA1 and HD mice; and X.W.Y helped interpret the results. P.G. organized data and figures for the manuscript.

## Supplementary Figures

**Figure S1.**
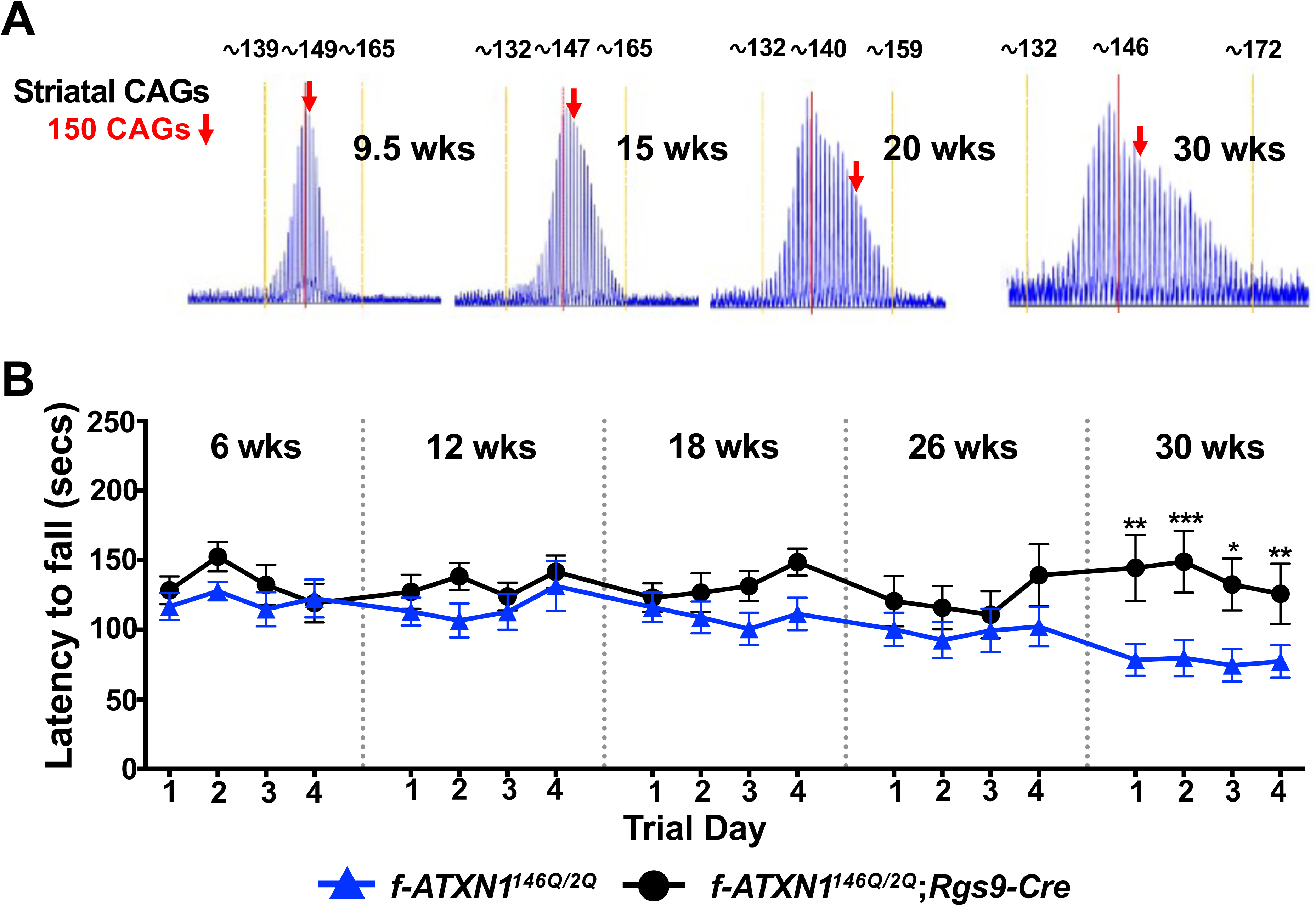
(Adapted from Fig. 3 of *Duvick et al., 2024*): Correlation of Striatal Somatic expansion with impact on motor function in *f-ATXN1^146Q/2Q^* mice. (**A**) The ABI GeneMapper trace plots showing the distribution of the PCR amplification products (peak height on the *Y*-axis and fragment size on the *X*-axis) generated with DNA isolated from the striatum of *f-ATXN1^146Q/2Q^* mice, at 4 different ages (in weeks: wks). The instability index value (number of CAGs) are shown on the graphs at each age. **(B)** Rotarod assessment for *f-ATXN1^146Q/2Q^;Rgs9-Cre*, and *f-ATXN1^146Q/2Q^* at 6, 12, 18, 26, and 30 weeks. Note the significant increase in the latency to fall at each trial day at 30 weeks when mutant ATXN1 is removed from MSNs (*f-ATXN1^146Q/2Q^;Rgs9-Cre*) compared to *f-ATXN1^146Q/2Q^*. Unpaired t test with Welch’s correction at each trial day for the given ages; *p=0.05; **p 0.01; ***p=0.001. See Supplementary Table 1 for complete statistics.

**Figure S2:**
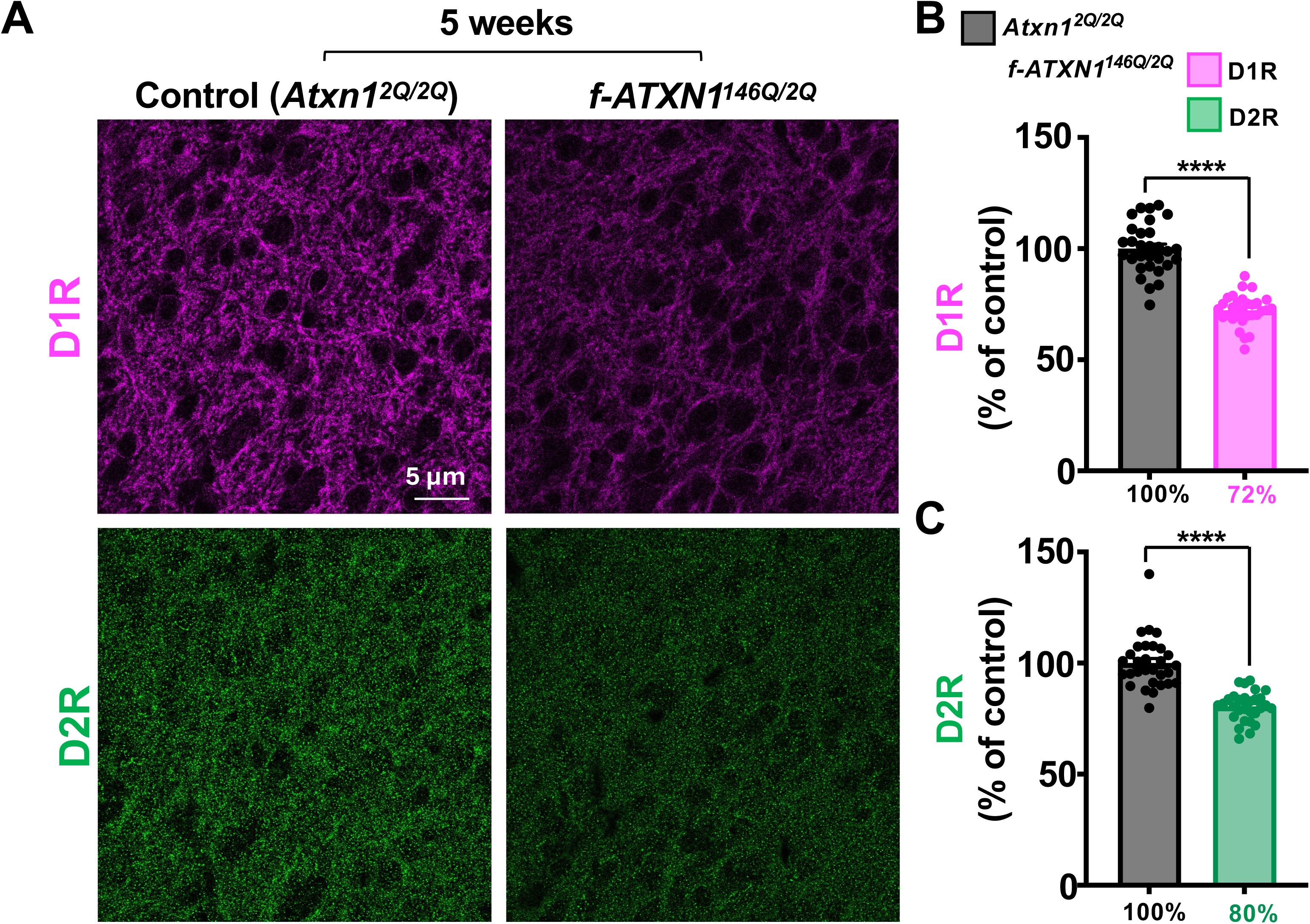
D1R and D2R protein expression is reduced in the nucleus accumbens of SCA1 mice. (**A**) Representative images of single sections of a z-stack acquired using confocal microscopy. Immunofluorescent staining using antibodies against D1R and D2R in the nucleus accumbens (NAc) of 5-week old *Atxn1^2Q/2Q^* (control) and *f-ATXN1^146Q/2Q^*(SCA1) mice. (**B-C**) A reduction in the fluorescence intensities of both D1Rs (**B**) and D2Rs (**C**) is observed in the NAc similar to the DLS (Fig. 2). Data is presented from 40-42 images across 3 mice for each genotype and normalized as percentage of average intensity of 40-42 images from 3 control mice. Unpaired t test with Welch’s correction; ****p=0.0001, ns=not significant, p>0.05. See Supplementary Table 1 for complete statistics.

**Figure S3:**
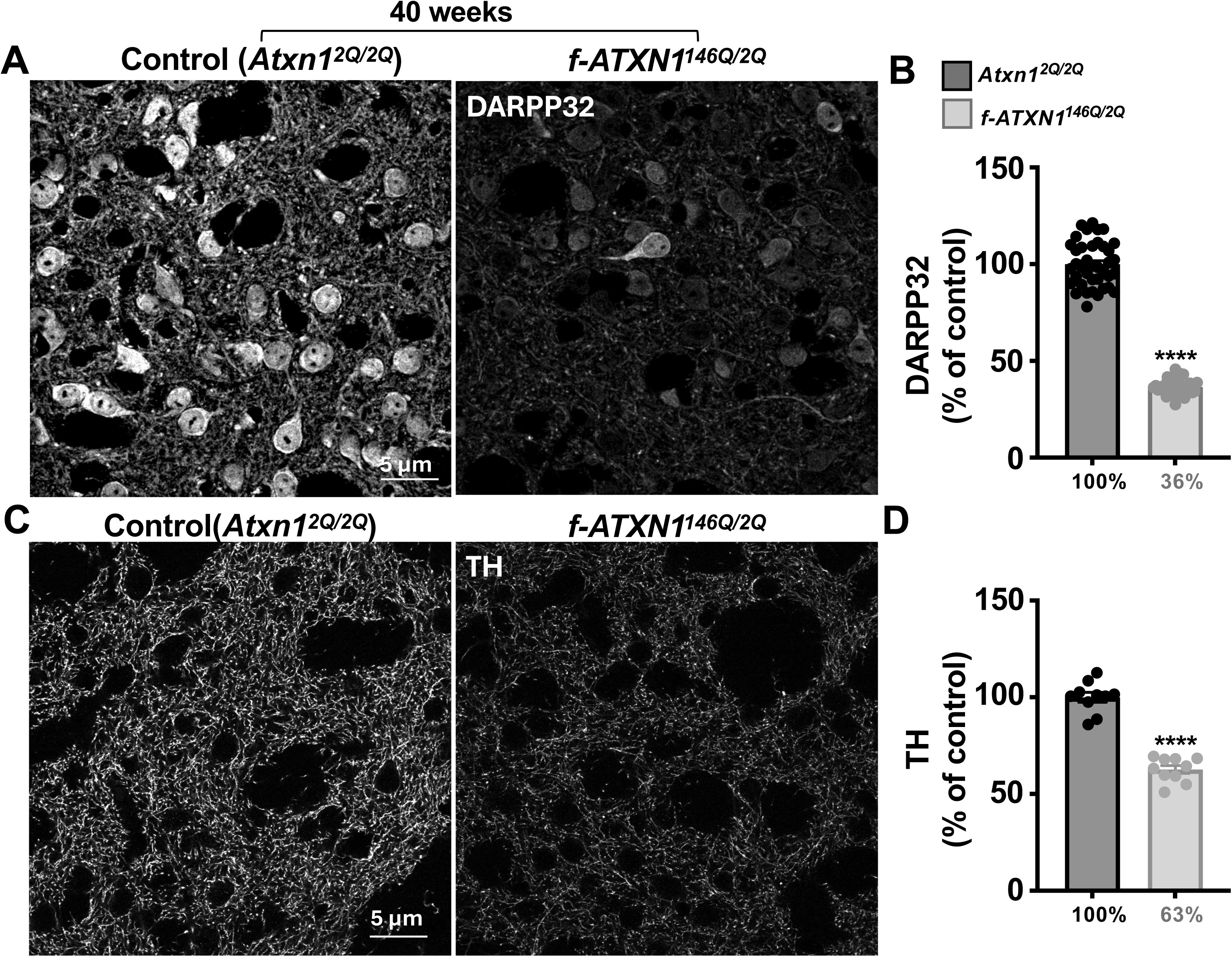
Expression of DARPP32 and tyrosine hydroxylase proteins is reduced in SCA1 mice. (**A-C**) Representative images of single sections of a z-stack acquired using confocal microscopy. Immunofluorescent staining using antibodies against DARPP32 (**A**) and tyrosine hydroxylase (TH; (**C**)) in the dorsolateral striatum (DLS) of 40-week old *Atxn1^2Q/2Q^* (control) and *f-ATXN1^146Q/2Q^*(SCA1) mice. (**B-D**) Quantification of reduced fluorescence intensity levels of DARPP32 (**B**) and TH protein (**D**) in SCA1 mice. Data is presented from 40-42 images across 3 mice for each genotype and normalized as percentage of average intensity of 40-42 images from 3 control mice. Unpaired t test with Welch’s correction; ****p=0.0001, ns=not significant, p>0.05. See Supplementary Table 1 for complete statistics.

**Figure S4:**
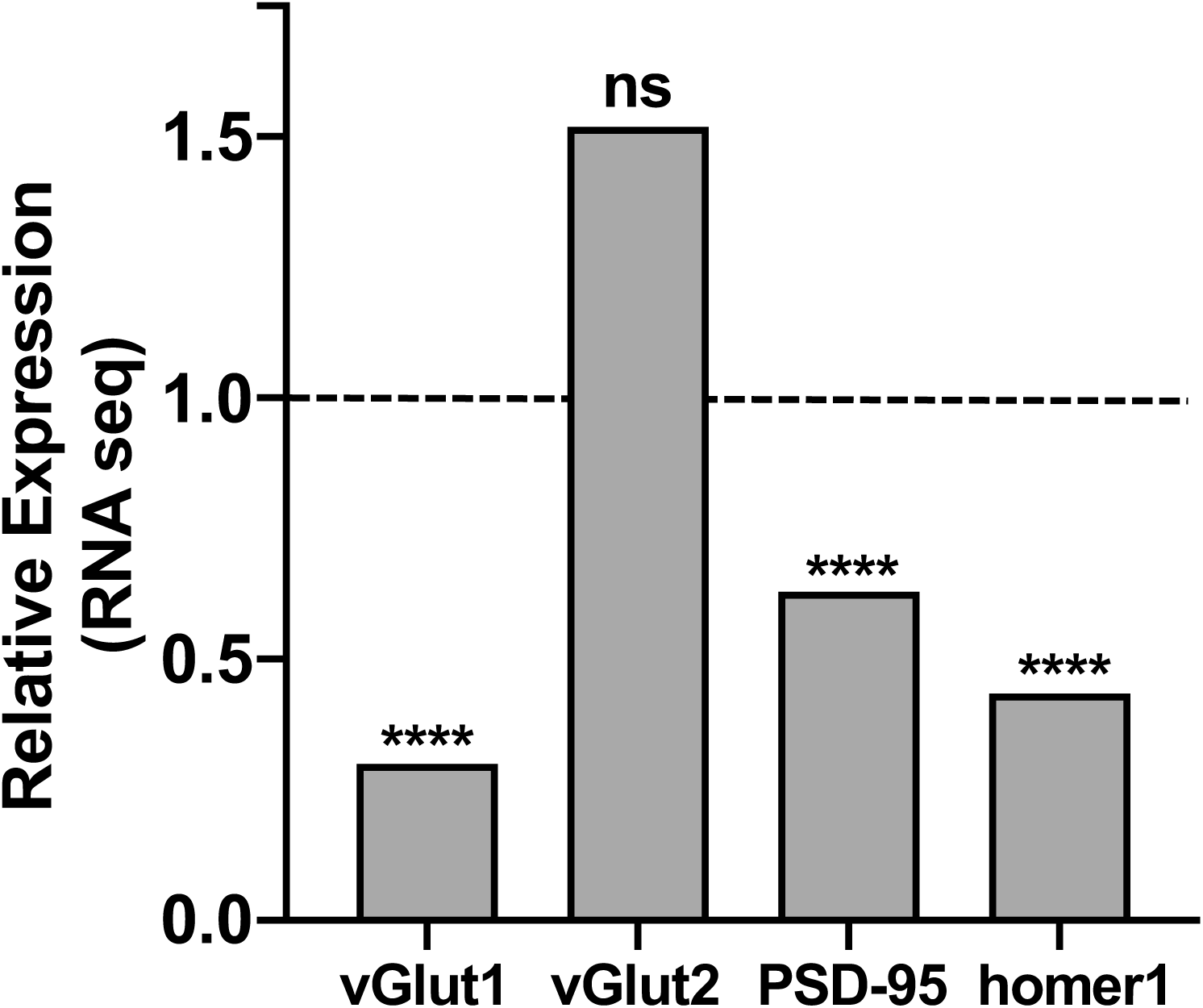
Transcripts of proteins at excitatory synapses are reduced in 10-week old SCA1 mice. Bar plots of gene expression as measured by RNA-seq in the striatum of 10-week old *Atxn1^175Q/2Q^* (SCA1) and control (*Atxn1^2Q/2Q^ ;*dashed black line) mice. Note the reduction in RNA levels of the presynaptic protein vGlut1 but not vGlut2, and a reduction in RNA levels of the postsynaptic proteins PSD-95 and homer1. Unpaired t test with Welch’s correction; ****p=0.0001, ns=not significant, p>0.05. See Supplementary Table 1 for complete statistics.

