## Supplementary Figures for "Striatal pathology in Spinocerebellar Ataxia Type 1 mice: A comparative study with Huntington’s disease"

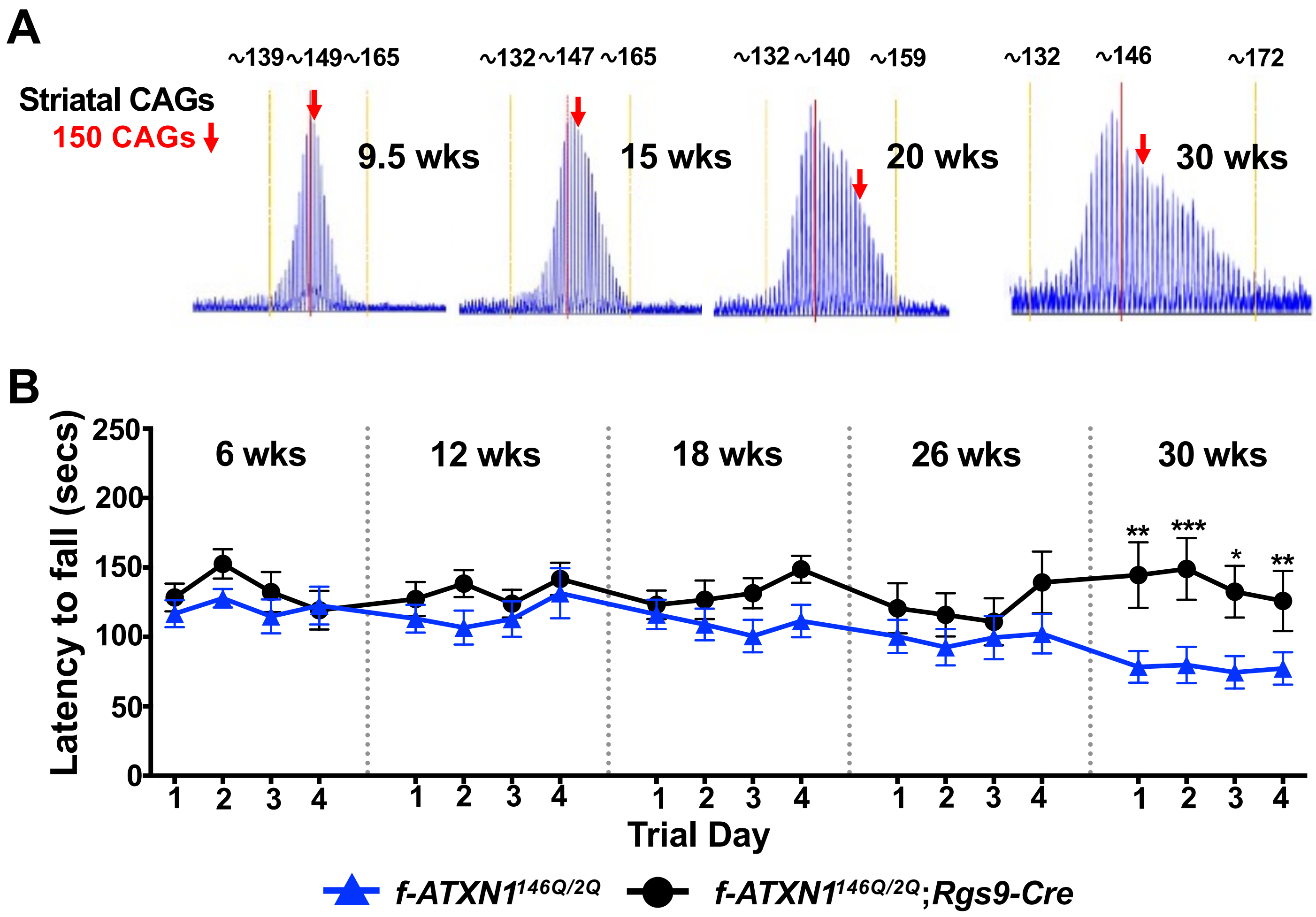


**Supplementary Figure 1**

**Figure S1 (Adapted from Fig. 3 of *Duvick et al., 2024*): Correlation of Striatal Somatic expansion with impact on motor function in *f-ATXN1*$^{146Q/2Q}$ mice.** (**A**) The ABI GeneMapper trace plots showing the distribution of the PCR amplification products (peak height on the *Y*-axis and fragment size on the *X*-axis) generated with DNA isolated from the striatum of *f-ATXN1*$^{146Q/2Q}$ mice, at 4 different ages (in weeks: wks). The instability index value (number of CAGs) are shown on the graphs at each age. **(B)** Rotarod assessment for *f-ATXN1*$^{146Q/2Q}$*;Rgs9-Cre*, and *f-ATXN1*$^{146Q/2Q}$ at 6, 12, 18, 26, and 30 weeks. Note the significant increase in the latency to fall at each trial day at 30 weeks when mutant ATXN1 is removed from MSNs (*f-ATXN1*$^{146Q/2Q}$*;Rgs9-Cre*) compared to *f-ATXN1*$^{146Q/2Q}$. Unpaired t test with Welch's correction at each trial day for the given ages; *p=0.05; **p 0.01; ***p=0.001. See Supplementary Table 1 for complete statistics.

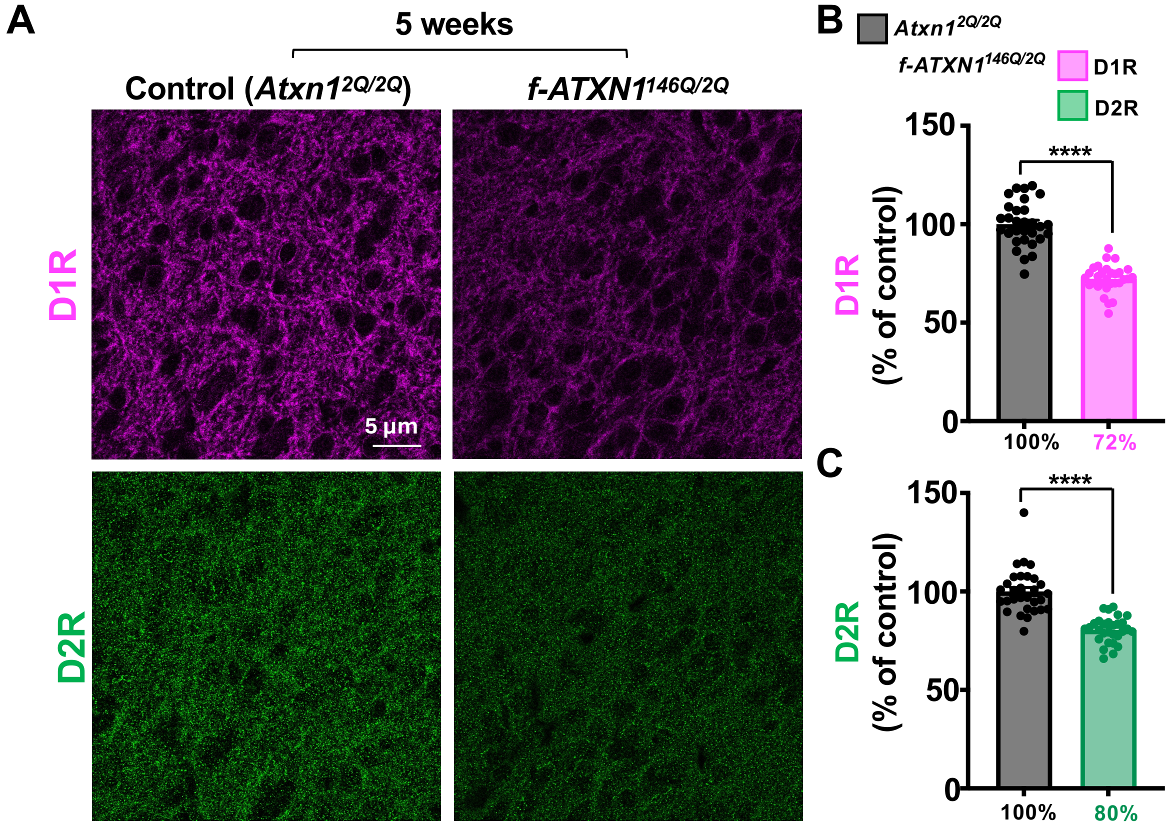


Supplementary Figure 2

**Figure S2: D1R and D2R protein expression is reduced in the nucleus accumbens of SCA1 mice.** (**A**) Representative images of single sections of a z-stack acquired using confocal microscopy. Immunofluorescent staining using antibodies against D1R and D2R in the nucleus accumbens (NAc) of 5-week old *Atxn1*$^{2Q/2Q}$ (control) and *f-ATXN1*$^{146Q/2Q}$ (SCA1) mice. (**B-C**) A reduction in the fluorescence intensities of both D1Rs (**B**) and D2Rs (**C**) is observed in the NAc similar to the DLS (Fig. 2). Data is presented from 40-42 images across 3 mice for each genotype and normalized as percentage of average intensity of 40-42 images from 3 control mice. Unpaired t test with Welch's correction; ****p=0.0001, ns=not significant, p>0.05. See Supplementary Table 1 for complete statistics.

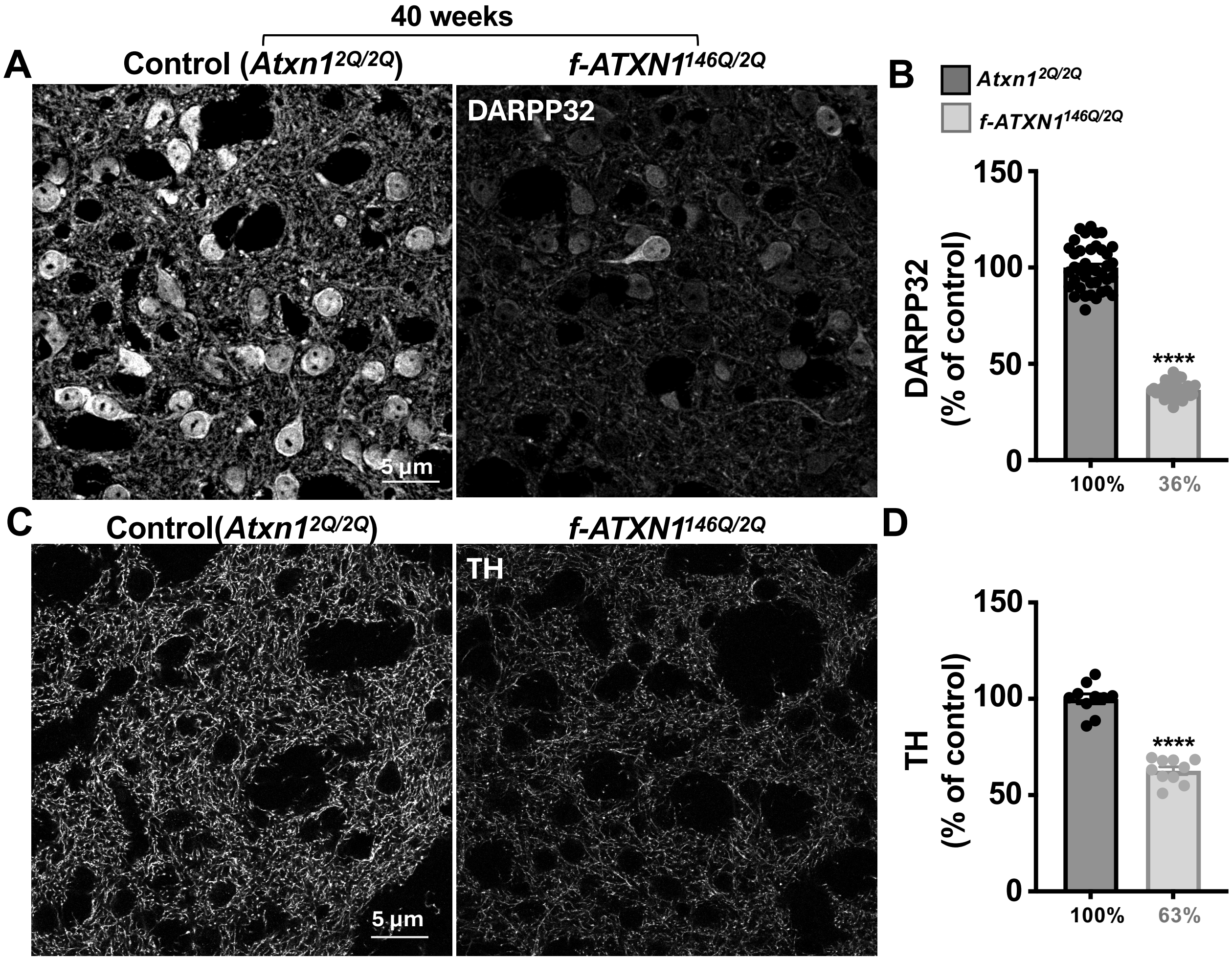


### Supplementary Figure 3

**Figure S3: Expression of DARPP32 and tyrosine hydroxylase proteins is reduced in SCA1 mice.** (**A-C**) Representative images of single sections of a z-stack acquired using confocal microscopy. Immunofluorescent staining using antibodies against DARPP32 (**A**) and tyrosine hydroxylase (TH; (**C**)) in the dorsolateral striatum (DLS) of 40-week old *Atxn1*$^{2Q/2Q}$ (control) and *f-ATXN1*$^{146Q/2Q}$ (SCA1) mice. (**B-D**) Quantification of reduced fluorescence intensity levels of DARPP32 (**B**) and TH protein (**D**) in SCA1 mice. Data is presented from 40-42 images across 3 mice for each genotype and normalized as percentage of average intensity of 40-42 images from 3 control mice. Unpaired t test with Welch's correction; ****p=0.0001, ns=not significant, p>0.05. See Supplementary Table 1 for complete statistics.

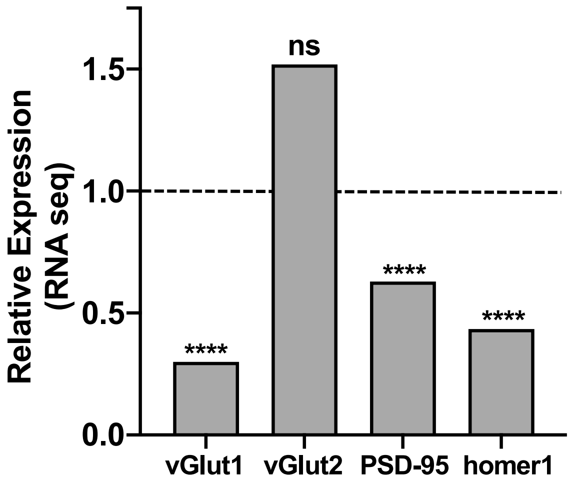


Supplementary Figure 4

**Figure S4: Transcripts of proteins at excitatory synapses are reduced in 10-week old SCA1 mice.**

Bar plots of gene expression as measured by RNA-seq in the striatum of 10-week old *Atxn1*$^{175Q/2Q}$ (SCA1) and control (*Atxn1*$^{2Q/2Q}$ *;*dashed black line) mice. Note the reduction in RNA levels of the presynaptic protein vGlut1 but not vGlut2, and a reduction in RNA levels of the postsynaptic proteins PSD-95 and homer1. Unpaired t test with Welch's correction; ****p=0.0001, ns=not significant, p>0.05. See Supplementary Table 1 for complete statistics.
